# Revision and annotation of DNA barcode records for marine invertebrates: report of the 8^th^ iBOL conference hackathon

**DOI:** 10.1101/2021.03.07.434272

**Authors:** Adriana E. Radulovici, Pedro E. Vieira, Sofia Duarte, Marcos A. L. Teixeira, Luisa M. S. Borges, Bruce Deagle, Sanna Majaneva, Niamh Redmond, Jessica A. Schultz, Filipe O. Costa

## Abstract

The accuracy of the identification of unknown specimens using DNA barcoding and metabarcoding relies on reference libraries containing records with reliable taxonomy and sequence quality. A rampant growth in barcode data led to a stringent need for data curation, especially in taxonomically difficult groups such as marine invertebrates. A major effort in curating marine barcode data deposited in the Barcode of Life Data Systems (BOLD) has been undertaken during the 8^th^ International Barcode of Life Conference (Trondheim, Norway, 2019). For practical reasons, only major taxonomic groups were reviewed and annotated (crustaceans, echinoderms, molluscs, and polychaetes). The congruence of Linnean names with Barcode Index Numbers (BINs) was investigated, and the records deemed uncertain were annotated with four tags: a) MIS-ID (misidentified, mislabeled or contaminated records), b) AMBIG (ambiguous records unresolved with the current data), c) COMPLEX (species occurring in multiple BINs), and d) SHARE (barcodes shared between species). A total of 83,712 specimen records corresponding to 7,576 species were reviewed and 39% of the species were tagged (7% MIS-ID, 17% AMBIG, 14% COMPLEX, and 1% SHARE). High percentages (>50%) of AMBIG tags were recorded in gastropods, whereas COMPLEX tags dominated in crustaceans and polychaetes. This high proportion of tagged species reflects either flaws in the barcoding workflow (e.g., misidentification, cross -contamination) or taxonomic difficulties (e.g., synonyms, undescribed species). Although data curation is crucial for barcode applications, such manual efforts of reviewing large datasets are not sustainable and the implementation of automated solutions to the furthest possible extent is hi ghly desirable.

## Introduction

Reference libraries, comprising sets of compliant DNA sequences assigned to species, are the backbone of species identification systems based on DNA barcoding and metabarcoding and thus, a crucial component in molecular biomonitoring (Weigand et al. 2019). Over the past 15 years, reference libraries experienced a rampant growth in the available number of DNA sequences and represented species (Porter and Hajibabaei 2018). To date, the ever -growing libraries have been deposited mostly in two large and public molecular databases, namely (i) GenBank (Sayers et al. 2021), a repository with data usually released after publication, and (ii) the Barcode of Life Data Systems (BOLD, Ratnasingham and Hebert 2007), a workbench in which data can be validated and analyzed before being released.

Parallel with this growth, instances of inaccurate or discordant data have been increasingly reported (Lis et al. 2016, Meiklejohn et al. 2019, Ramirez et al. 2020, Fontes et al. 2021), particularly when comparing data of morphological and molecular origin. This is recognized as a pertinent issue for the reliability of current and prospective DNA-based biomonitoring (Leese et al. 2018). Whereas any data discordances or inaccuracies in reference libraries would be more apparent using the “conventional” low-throughput DNA barcoding, they can be easily overlooked when applying high-throughput metabarcoding, where automated bioinformatics tools assign taxonomy to millions of sequences (Cristescu 2014). Because taxonomic assignments for metabarcoding data often do not undergo further scrutiny, any faulty assignments due to inaccurate reference library data can be repeated over and over without being noticed (Leese et al. 2016). The presence of errors in the reference libraries for a particular taxon (i.e. incorrect sequence assigned to a species) can nullify legitimate sequences since contradictory results cannot generally be managed by algorithms. The potential implications of this misleading data for biomonitoring and ecological assessment could be significant, particularly in the long -term and with the envisaged implementation of semi -automated and broad-scale metabarcoding-based biomonitoring systems (e.g., BIOSCAN, Hobern 2020).

Data discordances in reference libraries have multiple origins that can be split into two broad categories. First, there are discordances that are due to real biological complexities. Some of these are merely a reflection of the inherent uncertainties and dynamics of alpha taxonomy (e.g., Padial and De La Riva 2006, Padial and De la Riva 2020). Others originate from a mismatch between morphological and molecular diagnosis of species boundaries, that is, species that, for intrinsic reasons, can be diagnosed morphologically but not through short DNA sequences, or reciprocally. The second category of discordances are those introduced through operational errors. For instance, morphology -based specimen misidentifications have been often reported as a source of error (Pentinsaari et al. 2020). The experience level of the identifier may play a role in such errors, but often taxonomic keys, drawings and descriptions are incomplete or have poor quality, contributing to misidentifications. This may happen for some species that are frequently, but wrongly, recorded across the globe as cosmopolitan species for the simple reason that the original description is vague enough to accommodate various other species that remain undetected (Gómez et al. 2007, Padial and De la Riva 2020). Taxonomic variants such as synonyms and alternate representation designating the same taxon, are an additional source of mismatches (e.g., *Magallana gigas* and its alternate representation, but widely used name, *Crassostrea gigas*, Salvi et al. 2014, Bayne et al. 2017, Backeljau 2018). These types of inaccuracies and limitations are customarily shared and experienced by biodiversity databases (Bidartondo 2008, Patterson et al. 2010, Meiklejohn et al. 2019). Data discordances due to operational errors are also known to arise during collection, sampling and laboratory procedures, such as specimen and/or tissue sample mislabeling, cross -contamination, or non-targeted PCR amplification (Buhay 2009, Siddall et al. 2009, Evans and Paulay 2012). Poor sequence quality, sequencing errors or usage of reverse strand sequences may also contribute to discordances (reviewed by Pentinsaari et al. 2020).

A workbench such as BOLD, where data can be easily corrected if needed, brings great value to the barcoding and metabarcoding pipelines. Several tools for automated data quality control have been implemented in BOLD, including flags to indicate if sequences (COI, MatK, RbcL, RbcLa, trnH-psbA, ITS, ITS2) are barcode compliant or if they include stop-codons or common contaminants (e.g., human, cow, mouse, pig, bacteria). In addition, a number of analytical tools allow data congruence verification. For instance, discordances between species names assigned to records by BOLD users and the Barcode Index Numbers (BINs, Ratnasingham and Hebert 2013), automatically assigned to COI sequences by BOLD, can be easily revealed through the BIN discordance report. However, while automated bioinformatic routines can easily implement these flags and verification tools (e.g., Andújar et al. 2021), other inconsistencies ultimately require human-mediated inspection and judgement. In fact, BOLD staff review data and signal records for exclusion from the identification engine (BOLD IDS) if a potential misidentification or contamination has been detected. However, the sheer volume and diversity of data to be handled is not amenable for a comprehensive review and would require a more structured and feasible approach.

Reference libraries have been populated in part through dispersed contributions, in spite of a few central core facilities providing major inputs (e.g., Canadian Centre for DNA Barcoding). As a result, DNA sequence data, and respective metadata, which are uploaded to genetic data repositories such as BOLD or GenBank, have varied components and levels of compliance. The research practice also differs among target taxonomic groups, affecting even the type of vouchering system and metadata typically collected and accompanying each specimen (Rimet et al. 2021). The diversity of contributors and the idiosyncrasies of their research practices offer opportunities for operational discordances and shortcomings, eventually leading to increased difficulty in revision and curation of reference libraries.

The case for reviewing barcode data for marine invertebrates is particularly relevant. Marine invertebrates are often studied as a community and are one of the customary targets for marine biomonitoring, through metabarcoding (Duarte et al. 2021). The reference libraries are still far from complete and just beginning for some groups (McGee et al. 2019, Weigand et al. 2019, Leite et al. 2020) or geographical regions. In some cases, it has been estimated that several decades may be required to robustly populate the DNA barcode libraries (Vieira et al., in review). Beyond these difficulties, data inaccuracies or discordances occur with some frequency, possibly due to the more incipient status of the overall taxonomic knowledge and various other difficulties including the taxon-specific research practice and reduced number of available taxonomic experts, compared to other more well-studied groups such as vertebrates and insects (Radulovici et al. 2010, Mammola et al. 2020). Hence, marine invertebrates could greatly benefit from a comprehensive human-mediated revision and annotation of barcode data.

To accomplish this ambitious goal, of manually curating marine invertebrate barcode data, a hackathon was organized in the scope of the 8^th^ International Barcode of Life (iBOL) conference (Trondheim, Norway, 2019). A group of researchers involved in marine invertebrate barcoding were convened with the purpose of undertaking a comprehensive review and annotation of the barcode records of the most representative marine invertebrate taxa currently available in BOLD. This is a report on the approach, findings and implications for issues related to the curation of reference libraries of DNA barcodes.

## Methods

BOLD is a global database structured by few mandatory fields (e.g., phylum and country of collection), but habitat is an optional field and missing from many records, therefore a specific workflow (Fig. 1) was developed to consolidate only marine data for curation purposes. A copy of the World Register of Marine Species (WoRMS), the most comprehensive species list for the marine realm, was downloaded on June 1, 2019. Only accepted species names for extant marine animals were retained, and further filtering was performed to reduce the list to those invertebrate taxa that are mostly used in marine biomonitoring: crustaceans, echinoderms, molluscs, and polychaetes. For practical reasons to facilitate the curation process during the hackathon, only limited groups were selected in the final list: Crustacea (Malacostraca: Amphipoda, Isopoda, Mysida, and Euphausiacea), Echinodermata (all classes), Mollusca (Bivalvia and Gastropoda) and Polychaeta (all orders). The resulting subset of species from WoRMS was then compared against BOLD and matched species names had their BOLD public records added to pre-made BOLD datasets. Initially split by taxonomy (phylum), some of the larger datasets (e.g., molluscs) were further divided to reduce the number of records reviewed by one person during the hackathon. Only records with BIN assignments were retained. Within each dataset, a BIN discordance report was generated, as well as a neighbour-joining tree (NJ tree) with matching specimen details and images, if available, for each sequenced organism.

**Figure 1.**
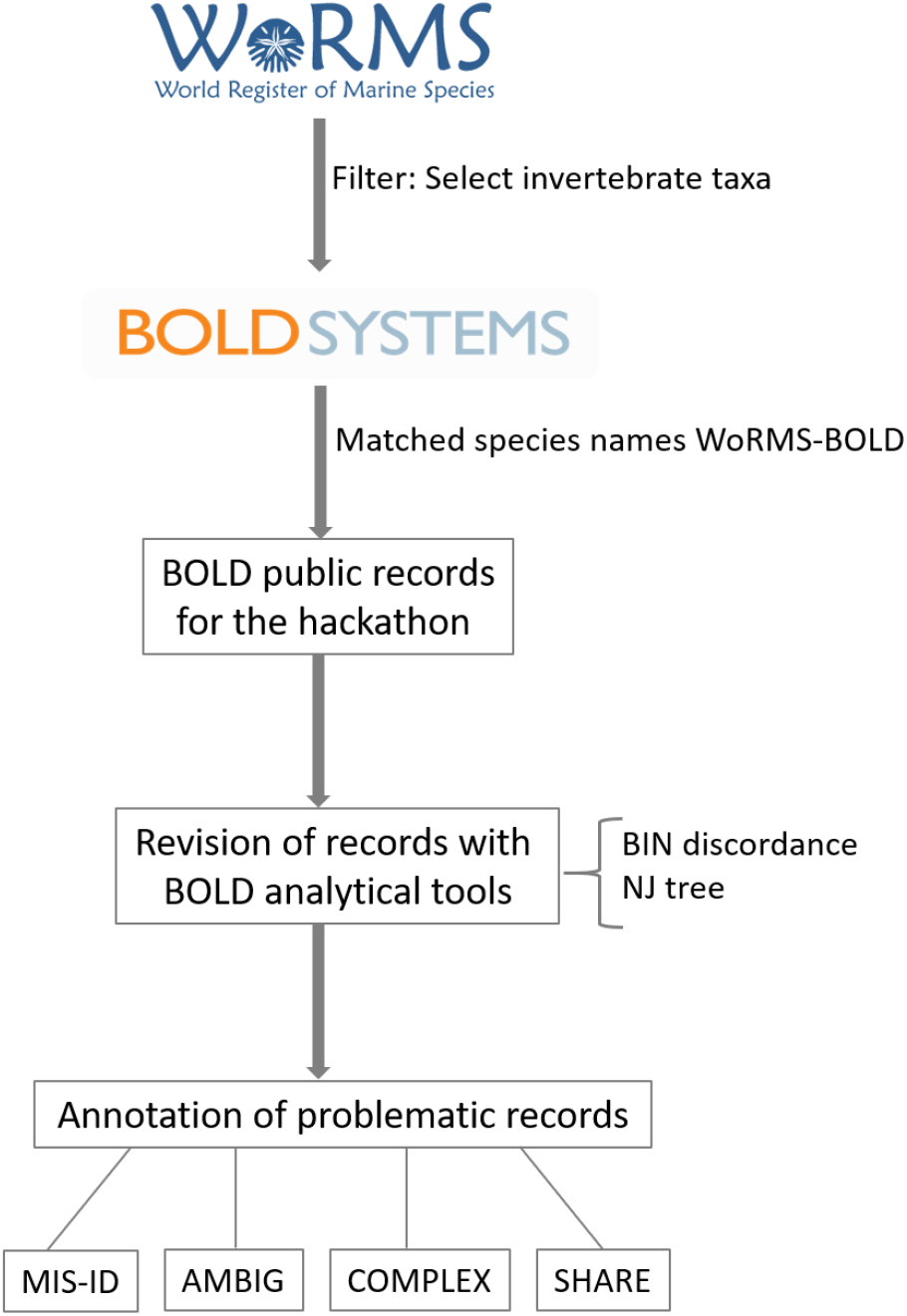
Workflow employed for review and annotation of select marine invertebrate records in BOLD. A subset of targeted invertebrate taxa was created from the initial list downloaded from WoRMS. This list was cross-referenced to the available taxonomic list from BOLD. Subsequently, only BOLD public records with BIN assignments for the matched species were placed in datasets and screened with some analytical tools (BIN discordance report, neighbour-joining (NJ) tree). Records deemed to be uncertain were annotated with four pre-established tags: MIS-ID (misidentification/mislabeling/contamination), AMBIG (ambiguous record), COMPLEX (species complex), SHARE (barcode sharing between species). Records suspected to be misidentified or contaminated were flagged by the BOLD team and removed from the identification system of BOLD.

The revision workflow (Fig. 1) consisted of manually inspecting the discordant BINs (i.e., multiple species names in one BIN) together with the NJ tree, followed by searching molecular databases (GenBank and BOLD) and published articles to gather additional information. Those records deemed uncertain based on the investigation conducted, were annotated with one of four pre-established tags:

a. MIS-ID (misidentification or contamination) – records believed to be misidentified, mislabeled or contaminated,
b. AMBIG (ambiguous) – records that could not be resolved with the current data,
c. COMPLEX (species complex) – records belonging to species with multiple BINs and, therefore, indicative of hidden or undescribed diversity,
d. SHARE (shared barcodes) – records belonging to species known to be sharing barcodes, due to incomplete lineage sorting or hybridization, based on existing literature.

All the records tagged as MIS-ID were submitted to the BOLD team so they can be flagged and, subsequently, removed from the database used for BOLD IDS. While detailed attention was given to discordant BINs, records in concordant BINs and singleton BINs were also reviewed, especially in cases of species with multiple concordant BINs (COMPLEX tag). The review of records (i.e., assignment of tags as well as additional notes) was recorded directly in the spreadsheets generated by BOLD as matching files for the NJ trees. Formulas were inserted to summarize the findings (number of records tagged, number of records and species per tag type, and number of taxa reviewed at each taxonomic rank). Due to the large amount of data requiring verification and the short time available, the work initiated during the hackathon continued during the months following the event. The results were illustrated through bar graphs using GraphPad Prism 9.0 (San Diego, CA, USA).

All the records reviewed can be found in BOLD (DS-HACK2019, DS-MOLL2019), while all the files with annotations are in the Supplementary Material.

## Results

The initial WoRMS download included ca. 600,000 taxa names at all taxonomic levels but only ca. 200,000 names were animal species with accepted names. Further filtering to invertebrate taxa of interest reduced the species list to 79,251 names as follows: Crustacea – 15,148 species, Echinodermata – 7,404 species, Mollusca – 44,883 species, and Polychaeta – 11,816 species. Only a fraction (about 10%) of these species had barcode representation in BOLD (Table 1).

Globally, the hackathon effort resulted in the review of over 83,712 DNA barcode records, distributed across 8,465 BINs, that corresponded to 7,576 marine invertebrate species from four phyla, 115 orders, 595 families and 2,490 genera (Table 1). Mollusca was by far the largest phylum tackled during the hackathon (ca. 50,000 records), therefore it was divided into two separate groups, Bivalvia and Gastropoda, for all the subsequent analyses presented here.

**Table 1.** Taxonomic classification and distribution of the reviewed DNA barcode records for the major taxonomic groups analyzed, and number of DNA barcode records and species that were tagged (MIS-ID, AMBIG, COMPLEX, SHARE) during the review of the dataset.

| **Taxonomic group** | **Phyla** | **Orders** | **Families** | **Genera** | **Species** | **BINs** | **DNA barcode records** | **Tagged species** | **Tagged records** |
| --- | --- | --- | --- | --- | --- | --- | --- | --- | --- |
| Bivalvia | Mollusca | 26 | 71 | 279 | 741 | 672 | 10,194 | 330 | 5,088 |
| Gastropoda | Mollusca | 38 | 233 | 1,066 | 3,982 | 4,235 | 39,749 | 1,582 | 15,581 |
| Crustacea | Arthropoda | 5 | 107 | 349 | 828 | 1,129 | 12,647 | 290 | 6,443 |
| Echinodermata | Echinodermata | 34 | 123 | 447 | 1,053 | 1,228 | 12,756 | 390 | 6,155 |
| Polychaeta | Annelida | 12 | 61 | 349 | 972 | 1201 | 8,366 | 365 | 3,434 |
| **Total** | **4** | **115** | **595** | **2,490** | **7,576** | **8,465** | **83,712** | **2,957** | **36,701** |

Gastropoda was the taxonomic group with the highest number of reviewed records (ca. 47%) and the highest number of species (53%) in the dataset (Table 1, Fig. 2). On the other hand, Polychaeta was the taxonomic group with the lowest number of reviewed records (ca. 10%), while Bivalvia was the group displaying the lowest number of species, comprising about 10% of the total number of species in the dataset (Table 1, Fig. 2).

**Figure 2.**
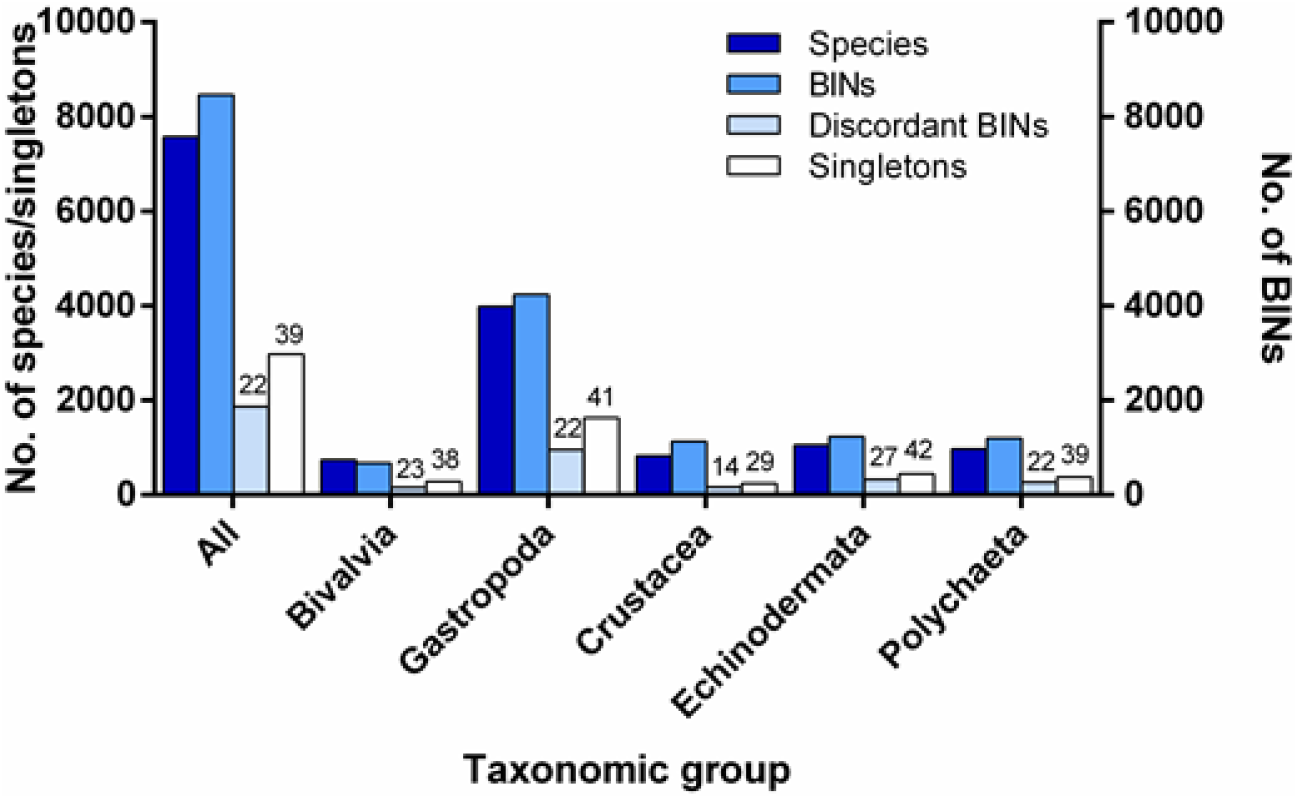
Number of species, BINs, discordant BINs, and singletons (species with only one DNA barcode record) for all groups analyzed and for each major taxonomic group separately. Numbers above bars indicate the proportion of discordant BINs and singletons, respectively.

The number of BINs was highest for Gastropoda and lowest for Bivalvia (Table 1, Fig. 2), and the ratio of BINs/species varied between 0.9 (Bivalvia) and 1.4 (Crustacea), suggesting that within most groups, some species displayed multiple BINs (Table 1, Fig. 2).

Across the entire dataset reviewed, ca. 22% of BINs displayed discordance (Fig. 2). The highest number of discordant BINs was found within the Gastropoda, while the lowest was within the Bivalvia. In terms of relative percentage, the Echinodermata was the group displaying the highest discordance (ca. 27%) and Crustacea the lowest (ca. 14%), in their BINs (Fig. 2). About 39% of the total number of analyzed species were singletons, i.e., represented by a single DNA barcode record on BOLD (Fig. 2). Echinodermata and Gastropoda were the taxonomic groups represented by a higher percentage of singletons (41 and 42%,respectively), while Crustacea the one represented by the lowest percentage (29%) (Fig. 2).

Nearly 39% of all species in the dataset were deemed uncertain (Fig. 3) and tagged with one of the four initially defined tags: MIS-ID (7%), AMBIG (17%), COMPLEX (14%) and SHARE (1%). Gastropoda had the highest proportion of tagged species (ca. 54%) and Crustacea the lowest (ca. 10%). Generally, most species were tagged with AMBIG (ca. 44%), and this ranged from ca. 30% to 50%, for Crustacea and Gastropoda, respectively (Fig. 4A). About 35% of the total tagged species were COMPLEX, with a higher percentage found among the Polychaeta (59%), Crustacea (ca. 58%) and Echinodermata (43%) and the lowest within the Bivalvia and Gastropoda (ca. 25% for each class) (Fig. 4A). On the other hand, only about 18% and 2% of the tagged species were classified as MIS-ID and SHARE, respectively (Fig. 4A). The highest proportion of the MIS-ID tag was found within the Bivalvia (ca. 29%), while the lowest was within Polychaeta (9%). Concerning the SHARE tag, the highest percentage was found within Gastropoda (ca. 4%), while Echinodermata and Polychaeta had no species tagged with SHARE (Fig. 4A).

**Figure 3.**
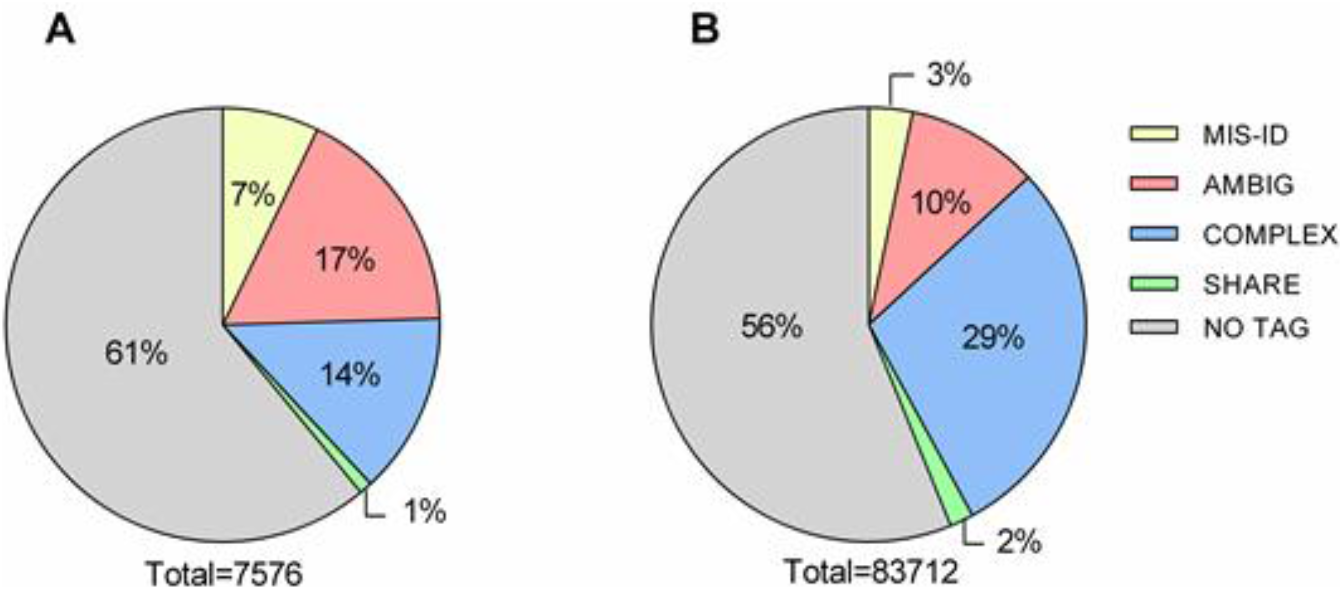
Distribution of the proportion of different tags in the reviewed dataset, in terms of species (A) and DNA barcode records (B). The total number of species (A) and records (B) are added below the chart.

**Figure 4.**
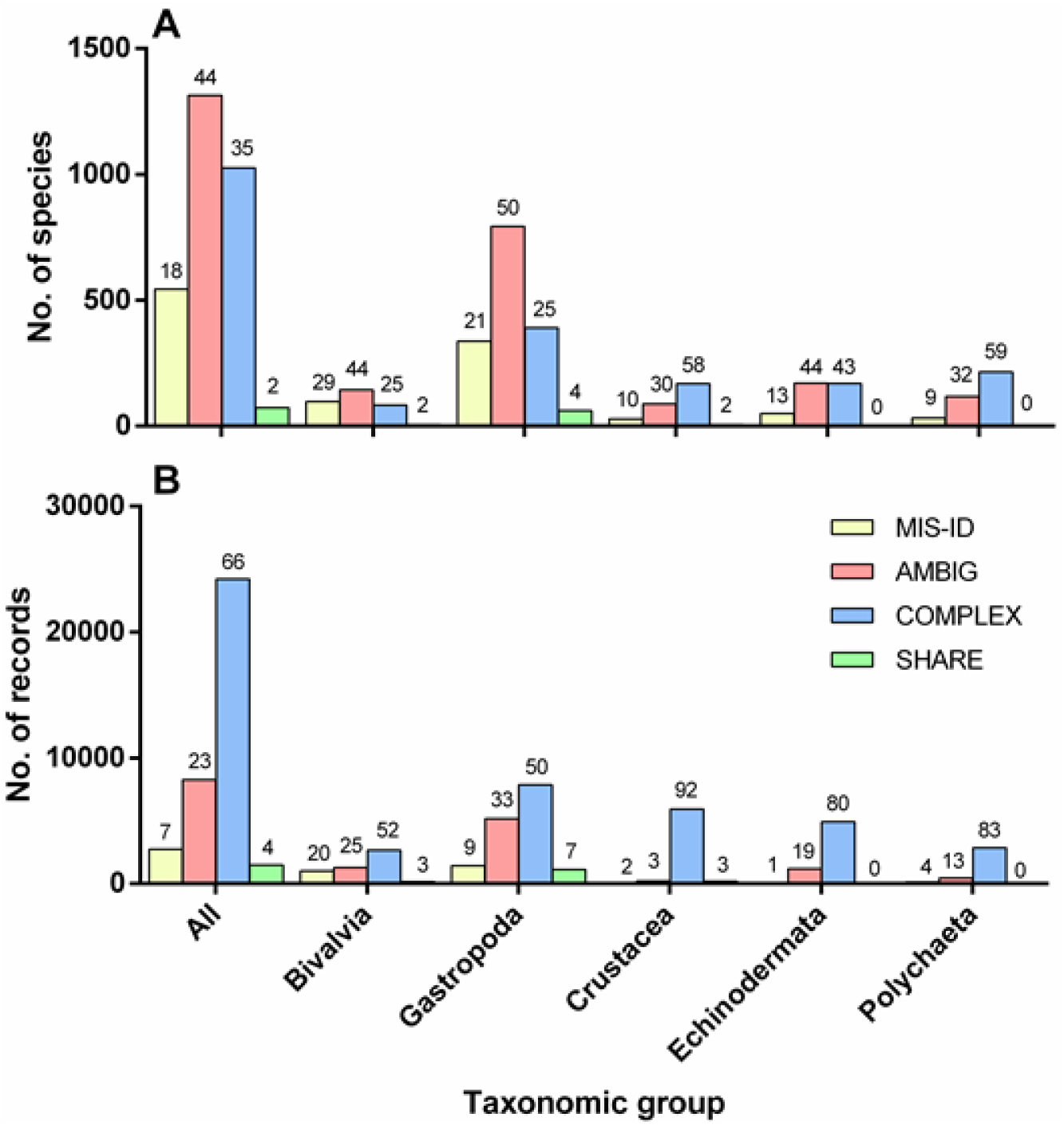
Distribution of the total number of species (A) and the total number of DNA barcode records (B), within each major taxonomic group, by the different tags. Numbers above bars indicate the relative proportion found within the whole tagged dataset and within each major taxonomic group.

Approximately 44% of all reviewed DNA barcode records were tagged with one of the four initially defined tags: MIS-ID (3%), AMBIG (10%), COMPLEX (29%) and SHARE (2%) (Table 1, Fig. 4B), with Gastropoda displaying the highest proportion (ca. 42%) and Polychaeta the lowest percentage of tagged records (ca. 9%). Generally, most records were tagged with COMPLEX (ca. 66%), and this varied between ca. 50% and 92% for Gastropoda and Crustacea, respectively (Fig. 4B). About 23% of the total tagged records were AMBIG, with the highest proportion found among the Gastropoda (ca. 33%) and Bivalvia (25%) and the lowest within the Crustacea (ca. 3%) (Fig. 4B). On the other hand, only about 7% and 4% of the tagged records were classified as MIS-ID and SHARE, respectively (Fig. 4B). The highest percentage of the MIS-ID tag was found within the Bivalvia (ca. 20%), while the lowest was within echinoderms (1%). Concerning the SHARE tag, the highest proportion was found within Gastropoda (ca. 7%), while no records were tagged with SHARE for Echinodermata or Polychaeta (Fig. 4B).

## Discussion

The hackathon on marine invertebrate barcodes served various goals beyond the immediate verification of the congruence between morphology and molecular data, and subsequent revision and annotation of records submitted to BOLD. To our best knowledge it constituted the first initiative of its kind for invertebrates (but see similar efforts in other taxa, Nilsson et al. 2018), thereby serving as a pilot initiative for similar actions. It also provided a unique human-mediated comprehensive assessment of the taxonomic congruence status of publicly available barcode data in BOLD.

At first glance, the results obtained for all taxa pointed out a considerable proportion of species records that may lead to erroneous identifications, particularly those tagged as MIS-ID and AMBIG (24% of reviewed species) and a relatively high proportion of species harbouring undescribed intraspecific diversity (14% of species tagged as COMPLEX).

MIS-ID tags constitute a relatively small portion of the reviewed species (7%) although it is possible that some AMBIG records are in reality MIS-ID but could not be fully resolved with the information available (see also further below regarding AMBIG). The review uncovered substantial differences in the proportion of MIS-ID between taxonomic groups, with incidence percentages up to five to six times higher among Bivalvia and particular groups of Gastropoda suggesting the need for sustained efforts on auditing further these two groups. Other than these uncertain groups, MIS-ID tags were below 4%.Although it is not a particularly worrying portion of the records, misidentifications may still have detrimental consequences, depending on the taxa and context of the study where the data is used (e.g. detection of non-indigenous marine species, see Bortolus 2008), and efforts to detect them should continue. One of the main accomplishments of the hackathon is that over 500 species (>2,700 records) of marine invertebrates were flagged and removed from BOLD-IDS, preventing this type of errors from being propagated in further studies using exclusively a molecular identification system.

A majority of AMBIG tags overall was not unexpected as the review employed a conservative approach and this tag was a last option when no other tag could be confidently assigned to uncertain records. This may have inflated the number of AMBIG tags that would have been assigned to other categories had this cautionary approach not been employed, although it is not possible to ascertain to what extent. On the other hand, a detailed taxa-partitioned inspection of the AMBIG records unravels a highly unbalanced distribution, with some particular taxonomic groups like Nudibranchia, Littorinimorpha and Pulmonata contributing disproportionately to the global numbers of tagged species (26%, 31% and 54%, respectively; Fig. S1), whereas in some other groups the proportion of AMBIG tags was comparatively low and even supplanted by other categories (e.g., Crustacea and Polychaeta). Littorinimorpha and Nudibranchia are particularly speciose taxa (e.g., 6,479 and 2,429 species respectively, WoRMS, June 1, 2019) that constitute unsorted taxonomic riddles, which underwent numerous taxonomic revisions, and of which the high number of synonymies (e.g. *Littorina obtusata* (Linnaeus, 1758) with more than 50 synonyms in WoRMS) are a manifestation. Pulmonata is no longer a valid taxon in WoRMS but it is still considered an order in BOLD, hence its use here.

It is important but difficult to distinguish between misleading data that originates from faulty procedures along the barcoding workflow, and imprecise data that results from poor basal taxonomic knowledge, unsolved taxonomic conundrums, unrecognized synonyms or unstable taxonomic status of particular taxa. Some of the AMBIG tags may result from misidentifications, others may in fact simply reflect unsolved taxonomies, that once sorted out may turn up to display congruence between molecular and other data sources. Eventually, part of the molecular data may even be evidencing the “true” species boundaries currently masked by complex morphological traces. AMBIG tagged records should therefore be taken as a signal for caution in their use, unless the end-user can find additional information for their clarification. A potential solution would be to avoid species-level identification when using these tags, giving preference to higher rank assignments (although an error at these ranks cannot be excluded with certainty). Recognition of taxonomic groups which have large number of AMBIG tags could provide a focus for more detailed taxonomic work to clarify the status of various species.

The COMPLEX tag is the second most prevalent overall, but it is also the only one that does not necessarily preclude the accurate identification of specimens. It simply signals cases where possible undescribed intraspecific diversity was found. Occurrence of multiple and highly divergent intraspecific lineages have been abundantly and increasingly reported in diverse groups of marine invertebrates, suggesting the existence of considerable hidden diversity (e.g. Nygren and Pleijel 2011, Lobo et al. 2017, Nygren et al. 2018, Borges and Merckelbach 2018, Desiderato et al. 2019, Teixeira et al. 2020). Several studies employed additional markers, mitochondrial and nuclear, essentially confirming the patterns observed with the DNA barcodes (e.g. Hupało et al. 2019, Vieira et al. 2019). Contrary to observations made for other tags, the Crustacea (particularly Amphipoda) and Polychaeta were the groups with a higher and substantial proportion of species tagged with COMPLEX, even more than MIS-ID and SHARE (Fig. 4), although this appears to be a common occurrence across the examined taxa, given that most phyla and classes displayed at least close to 10% or higher. Curiously, the Gastropoda displayed comparatively low values with this tag. This may reflect in fact lower incidence of high-intraspecific divergences in this group but may also result in part from truly COMPLEX tags masked in the AMBIG category due to the high number of taxonomic discrepancies in the group.

Although not so critical for the accuracy of identifications, at least according to the current status of taxonomic knowledge, there are important aspects of the COMPLEX tag to consider. Most notably, it helps when perceiving the overall quantity of presumptive marine invertebrate species waiting verification and eventual consolidation and description. Failing to recognize this considerable amount of hidden diversity may be just as detrimental for bioassessment and monitoring as the MIS-ID or AMBIG cases (Bickford et al. 2007). In general, very little is known about potential biological and ecological differences among the highly divergent lineages, some of them confirmed species. Some of the species assigned with COMPLEX are prominent indicator species included in biotic indexes for bioassessment. For instance, *Hediste diversicolor, Eumida sanguinea, Corophium multisetosum*, and several *Gammarus* species are included in the AZTI Marine Biotic Index species list (Borja et al. 2000, https://ambi.azti.es/). Failing to recognize their hidden diversity may contribute to imprecisions and lower performance of the ecological status assessments.

A number of marine invertebrates thought to have cosmopolitan or wide distributions are being unraveled as complexes of multiple units with much narrower distributions (e.g. Gomez et al 2005, Borges and Merckelbach 2018, Teixeira et al. 2020). It is not uncommon that divergent lineages are geographically arranged in segregated groups, so that genetic markers can be used not only to identify the morphospecies but also regional or local lineages (e.g. Hupało et al. 2019, Vieira et al. 2019). An accurate perception of changes in species distributions and occurrence, particularly in the scope of global change-induced responses, requires detailed recognition of highly divergent intraspecific lineages. Those presenting narrower ranges are also especially relevant from a biodiversity conservation perspective (Bickford et al. 2007) and consideration of marine protected areas and other conservation actions.

Species and records tagged with SHARE are by far the lowest proportion globally and within each taxonomic group. Part of the SHARE tags correspond to cases of low interspecific divergence coupled with incomplete sorting and haplotype sharing, or even a result of hybridization and introgression (particular cases can be found in the annotation files in the Supplementary Material). Hence, these reflect instances where the COI barcode sequences are unable to discriminate species, rather than a reference library flaw resulting from faulty steps in the barcoding workflow. Yet, they may also be applied to cases of completely sorted and well-established species whose records fall within the same BIN. Either way, these results indicate that the occurrence of SHARE cases is minimal and can be promptly flagged, or, in the latter case, circumvented through the accumulation of records into the libraries and refinement of the BIN assignment for that particular group. Gastropoda was again the group with the highest incidence of SHARE tags, reinforcing the perception that greater research effort is needed for taxonomic clarification of marine members of this group.

The BIN discordance report generated in BOLD is a very useful validation tool which easily highlights uncertain cases in need of careful examination. However, concordant BINs are not exempt from misidentification, especially less represented BINs, with barcodes from one project or from multiple projects where BOLD users did not hold taxonomic information for their specimens and relied solely on the existing database which, if erroneous initially, could be largely propagated. In addition, singletons are very difficult, if not impossible, to verify. As the hackathon data included about 30-40% singletons for each taxonomic group investigated, we cannot exclude a larger proportion of the current marine data that might need to be tagged with one of the four labels discussed above. However, the difficulties encountered in reviewing barcode data, and especially marine barcode data, are a signal for the need of better practices when generating, analyzing, and publishing barcode data.

## Conclusions

The one-day hackathon and the following months of annotation effort provided a significant contribution to the curation of the BOLD DNA barcode reference libraries for major groups of marine invertebrates. Although several large taxonomic groups had to be excluded from the analyses, it still constituted a massive undertaking, resulting in the individual review and annotation of a very large number of records and species.

Ideally, this type of event should be repeated on a regular basis, accompanying the accumulation of new records in reference libraries. However, as a corollary of this enterprise, it was very evident that the immense effort required to complete this task cannot be underestimated, and that it could hardly be repeated in the same format.

Indeed, a much more practical approach is needed in future endeavours, and this pilot exercise provided some possible solutions to substantially simplify the review procedure. Simplification could start by reducing the effort needed for preparing the data before review.

This can now be accomplished, for example, through recently developed applications such as BAGS (Fontes et al. 2021). Secondly, and most importantly, attempting to develop intelligent systems that can screen out the most obvious discordances and misleading records, and dispense human-mediated verification. The results presented here indicate that a considerable fraction of the discordances could fall in this category. The outstanding capabilities of machine learning (ML) and artificial intelligence (AI) systems have been extensively demonstrated in various research fields (Davenport and Kalakota 2019) including in image-based species identification (e.g. Ärje et al. 2020, Høye et al. 2021) and in the analysis of metabarcoding data for ecological status assessment (e.g. Cordier et al. 2019, Frühe et al. 2020). Nonetheless, it was also evident from our results that there will always be a fraction of discordances that cannot be addressed through automated systems and will require human intervention to be properly annotated.

Therefore, whereas ML and AI-type of approaches may help to considerably reduce the number of records requiring review, turning hackathon-like initiatives into practical and feasible commitments, at the end of the line there will be the need for human-mediated verification at least, and hopefully, for a minor set of records. In this regard, DNA barcode reference libraries are no different than other biodiversity data, and, ideally, strategies for data curation through community involvement (see the community of WoRMS editors curating data) could be used as inspiration and transposed to the DNA barcoding practice.

## Supporting information

AnnotationFiles

## Acknowledgements

The hackathon was organized with financial support from DNAqua-Net in the scope of the 8^th^ International Barcode of Life Conference in Trondheim, Norway on 16 June 2019. We thank DNAqua-Net for the funding provided and the local conference organizers for all the logistical support received. We are grateful to Tyler Elliot and the rest of the BOLD team for their help with data queries and analytics. We also thank the hackathon participants for vibrant discussions during and after the event: Berry van der Hoorn, Katrine Konsghavn, Guy Paz, Mouna Rifi, Malin Strand, Anne Helene Tandberg, Adam Wall, and Endre Willassen. Marcos A. L. Teixeira was supported by a PhD grant from the Portuguese Foundation for Science and Technology (FCT I.P.) co-financed by ESF (SFRH/BD/131527/2017). Financial support granted by FCT to Sofia Duarte (CEECIND/00667/2017) is also acknowledged. Pedro Vieira was supported by the project NIS-DNA (PTDC/BIA-BMA/29754/2017), also funded by FCT.

