## Supplementary material for "Revision and annotation of DNA barcode records for marine invertebrates: report of the 8^th^ iBOL conference hackathon": Figure S1: Figure S1.docx

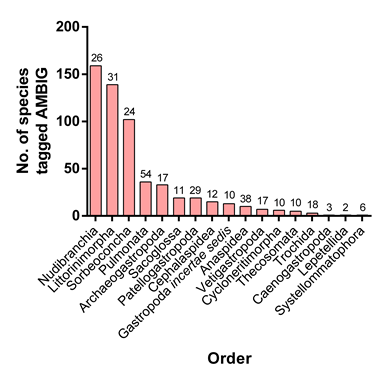

Figure S1. Number of species tagged with AMBIG within each order in the Gastropoda. Numbers above bars indicate the relative percentage of species tagged AMBIG, within each order. Only orders with species tagged AMBIG and with more than 10 species are displayed in the figure.
